# Integrating animal experience into state-switching habitat selection models

**DOI:** 10.64898/2026.07.28.741089

**Authors:** Romain Dejeante, Akina Kuperus, Mark. A Lewis, John M. Fryxell

**Author notes:** These authors contributed equally (Romain Dejeante, Akina Kuperus).

## Abstract

Movement models often assume that animal decisions depend on perceived environmental conditions. However, optimal foraging and cognition theory predict that accumulated experiences should drive changes in animal motivation and decision-making. Here, we propose a mechanistic framework that models state-switching habitat selection as a history-dependent process, where internal states, such as fear or energy, emerge as latent variables accumulating through environmental exposure. Simulations showed that our model accurately recovers memory timescales and cumulative effects of environmental exposure on behavioural switching that would be undetected by existing state-switching habitat selection models. Applied to woodland caribou, it reveals that an individual integrates predation risk experienced over the previous 15 days, but food intake over only 2 days, when deciding to remain or leave an area. Our model advances perspectives on movement ecology by quantifying how accumulated experiences shape changes in motivation driving animal movement decisions.

## 1. Introduction

Animal movement is often conceptualized as a complex interplay between an individual’s internal state, mechanical capacity for motion, navigational capacity, and fitness-influencing features of the surrounding environment, such as food availability, spatial familiarity, or predation risk (Nathan et al. 2008). Classical cognition theory generally formulates that spatially driven memory plays a central role in shaping movement decision-making (Fagan et al. 2013). A growing body of mechanistic movement models, parameterized with empirical data, demonstrated that animals inform their movement decisions by relying on spatial memory (i.e., memorizing previously visited locations) (Falcón-Cortés et al. 2021; Merkle et al. 2019; Rheault et al. 2021; Schlägel et al. 2017; Thompson, Lewis, et al. 2022) or attribute memory (i.e., memorizing the environmental features associated to these locations) (Avgar et al. 2015, 2013; Gurarie et al. 2022; Merkle et al. 2014; Ranc et al. 2021, 2022; Wolf et al. 2009). These models fundamentally quantify *where* an animal moves in regard to spatial or attribute memory (i.e., selecting or avoiding known places), for example by coupling standard movement models, such as step selection function (SSF), with cognitive maps (Kim et al. 2024). Less attention has been placed, however, on the potential role of previous experience, mediated by physiology or cognitive processes, in shaping movement decisions through its accumulative effect on motivational state.

Animals routinely experience different levels of risk of predation or rates of energetic flux when travelling through a landscape (Laundre et al. 2010; Shepard et al. 2013), influencing their fear state or their energetic condition which may, in turn, influence their movement decisions. These physiological or psychological states are commonly referred to as ‘internal states’ in movement ecology and describe the motivational states of an individual to satisfy an internal need (i.e., gaining energy, avoiding risk, reproducing) (Nathan et al. 2008). Changes in an animal’s internal state should motivate changes in animal decision-making. For example, optimal foraging and energy-safety theories formulate that the energetic state of an individual should influence its movement decisions. Optimal foraging theory suggests that individuals adopt ballistic movement to search for patchy resources when hungry and area-restricted searches to exploit resources once a rich feeding patch is found (Benhamou 1992; Dorfman et al. 2022; Kareiva and Odell 1987). From an energy-safety theory perspective, individuals should select foraging areas despite their elevated predation risk when their energetic reserves are low, but seek safety when they have accumulated enough energy to meet their future needs (Houston et al. 1993; McNamara and Houston 1992).

An animal’s internal state may depend not only on the immediate environmental conditions but also on the accumulation of experiences with its environment. For example, an individual’s energetic reserves at a given time are determined not only by current resource availability but also by the energetic gains and costs accumulated over time (Mangel and Clark 1988). Similarly, an individual’s fear level should depend on both short-term and long-term exposure to predation risk, rather than solely on the immediate perception of a predator (Creel 2018). Proactive responses to the risk of predation require long-term, predictable exposure to a predation risk and should rely on an animal’s memory of previously perceived risk (Creel 2018). The motivational states of an individual may therefore be a function of both immediate environmental conditions and previous individual experiences that can be stored through cognitive or physiological processes.

Hidden Markov models (HMM) provide an opportunity to explicitly model the dynamics in animal motivational states and, when coupled with step selection functions (HMM-SSF), allow inferences on state-dependent habitat selection (Klappstein et al. 2023; Pohle et al. 2024; Prima et al. 2022). Although the identified states are commonly referred to as ‘behavioural states’, they arise by discriminating several modes of habitat selection and/or movement properties, such as periods of high selection vs. periods of low selection for a given habitat type. However, although existing HMM-SSFs explicitly model the transition between distinct modes of habitat selection, state dynamics are currently modelled as a function of immediate environmental conditions rather than as a physiological or psychological process that accounts for an animal’s previous experiences. Incorporating cognitive maps into HMM-SSFs allows ecologists to account for the influence of spatial or attribute memory on habitat selection (Thompson, Derocher, et al. 2022; Thompson, Lewis, et al. 2022), but it does not capture how the cognitive process drives changes in animals’ motivational states. Addressing this limitation requires relaxing the null assumption that motivational states are independent of prior experiences, rather than simply adding an additional covariate to the step selection function. Yet, these HMM-SSF models are increasingly popular and broadly accessible to ecologists, contrary to other movement models accounting for animals’ internal states, such as chemotaxis models (Erban and Othmer 2004), prey-taxis models (Kareiva and Odell 1987), and continuous-time recharge-based movement model (Hooten et al. 2019).

Here, we propose a mechanistic framework to evaluate how memory-based motivational states influence movement decision-making. We introduce a memory-based hidden Markov model coupled with a step selection function that explicitly incorporates past experiences into the internal dynamics that govern transitions between states. Unlike existing memory-based movement models that rely on cognitive maps, our memory process models the impact of cumulative experiences on transition rates between behavioural states, relaxing the assumption that state transitions depend solely on current environmental conditions. We model transition probabilities as functions of temporally weighted past experiences and estimate the weight attributed to these experiences. In doing so, our model allows ecologists to test whether animals rely on the memory of past experiences to motivate their movement states, addressing the question of *why* animals move. Using simulated data, we demonstrate that memory-based state-switching habitat selection models can accurately estimate the weight attributed to previous experiences, as well as common estimates of these models (i.e., step selection coefficients and coefficients of transition between states), where standard HMM-SSF models would yield biased estimates for state-switching probabilities. When applied to woodland caribou (*Rangifer tarandus caribou)*, we demonstrate that an individual may rely on predation risk experienced over the previous 2 weeks to decide whether to stay or leave an area, rather than from the immediate risk of predation.

## 2 Methods

### 2.1 Model Formulation

To evaluate how animals’ previous experiences influence movement decision-making, we introduce a memory-based Hidden Markov model (memory-based HMM) coupled with a step selection function (SSF) in order to model simultaneously (1) the weight that animals attribute to previous experiences, (2) how temporally weighted experiences influence transition between modes of habitat selection and (3) the habitat selection coefficients that characterize these modes of behaviours.

#### 2.1.1 Memory-based hidden Markov model

We introduce a memory-based hidden Markov model to describe the changes in an animal’s internal state. The number of states is decided *a priori* (Pohle et al. 2017) and the HMM describes the probability of transitioning between states. This dynamical system is conditional first-order Markovian, meaning that transition probabilities depend only upon the current state, conditioned upon auxiliary variables that can include memory.

Let {*S*_1_, *S*_2_, …, *S*_*T*_} be the sequence of states of an animal (for example, exploratory or encamped states). For a *K*-state memory-based HMM, the probabilities of transitioning from one state to another at each time step *t* are given by the *K* × *K* transition matrix:

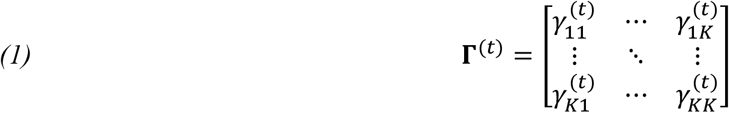

Where 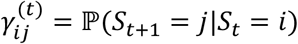, the probability of transitioning from state *i* to state *j*. Each state has associated movement and habitat selection parameters that describe the observable movement behaviour for that state (more on this in 2.1.2).

The transition probabilities are dependent on temporally weighted previous experiences of covariates (*ω*_1_, *ω*_2_, ⋯, *ω*_*P*_) . For example, covariates could include the risk of predation perceived at a given time, the amount of energy available at a given location, the presence of a social partner or competitor or a specific interaction associated with that individual (such as nursing, mating, or contest events), or the intensity of human disturbance experienced at a given location.

For each covariate,

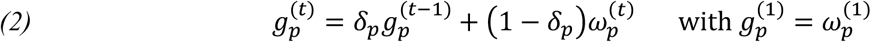

is the accumulated memory of that covariate. The parameter 0 ≤ *δ*_*p*_ ≤ 1 determines the proportion of weight given to past experiences of covariate ω_*p*_, with the remaining weight (1 − *δ*_*p*_) given to the current experience. A linear predictor function

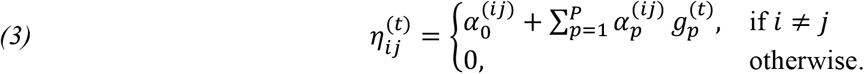

describes the effect of the accumulated memory on the transition probabilities for each time step via a multinomial logit link function:

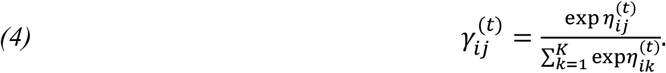

This memory-based HMM allows transition probabilities to depend on both the covariate experienced at time *t*, as well as all past experiences, with experiences further in the past given lower weights. In an animal movement context, 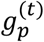 may reflect, for example, energetic states or fear levels. It is important to highlight that 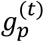 is not a spatial variable, but rather a measure of accumulated previous experiences that are hypothesized to influence state switching and movement behaviour. For example, if ω_*p*_ is the amount of resources available at a specific location or the risk of predation perceived at a given time, *g*_*p*_ reflects the accumulated energetic state or levels of fear experienced over a geometrically weighted time horizon into the past.

Notice that Equation [2] is recursive. At time *t*, the weight of the experience at time *t* − *ℓ* is (1 − *δ*)*δ*^*t*−*ℓ*^. Thus, the weight attributed to each previous experience is a monotonically decreasing function of time since the experience, reflecting the decay in ‘memory’ of past experiences (Appendix A. Figure S1). When the weight attributed to previous experiences is null (*δ*_*p*_ = 0), we recover the original HMM described in Klappstein et al. (2023) without memory.

#### 2.1.2 Step selection function

To incorporate movement behaviour into the model, we link the memory-based HMM with a step selection function (SSF) which describes the observable movement of an individual (Avgar et al. 2016). Let {*s*_1_, *s*_2_, …, *s*_*T*_} be the animal locations observed at regular time intervals. The SSF describes the likelihood of an animal being at location *s*_*t*_ at time *t*, given their previous locations *s*_*t*−1_, …, *s*_1_ (Fortin et al. 2005). The likelihood is dependent on both the animal’s movement capacities, described by the movement kernel *ϕ*, and their habitat selection, described by a weighting function *w*, and is given by

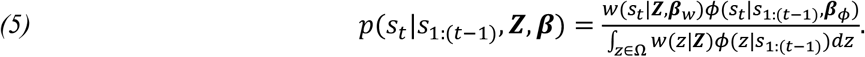

The integral in the denominator is taken over the study area Ω and ensures that Equation [5] is a probability density function with respect to *s*_*t*_. This integral is intractable but can be approximated using Monte Carlo integration methods (Michelot et al. 2024). The weighting function is a log-linear function of habitat covariates *Z* = (*Z*_1_, …, *Z*_*m*_) with selection parameters ***β***_*w*_ = (*β*_1_, …, *β*_*m*_). The choice of covariates used in the weighting function are assumed to impact the spatial movement decision making of an animal on the time scale at which the locations are observed.

Coupling the memory-based HMM with an SSF allows movement and habitat selection to depend on an animals’ state. For an animal in state *S*_*t*_ = *k* and location *s*_*t*_ at time *t*, the SSF for the next step would be *p*^(*k*)^(*s*_*t*+1_|*s*_*t*_, *s*_*t*−1_; ***β***^(*k*)^) where ***β***^(*k*)^ are the movement and habitat selection parameters associated with state *k*. With the memory-based HMM, the transition between states accounts for both current and past experiences.

### 2.2 Model fitting

For a sequence of observed locations {*s*_1_, *s*_2_, …, *s*_*T*_}, the likelihood of the memory-based HMM-SSF model (where the SSF is first-order Markovian) is

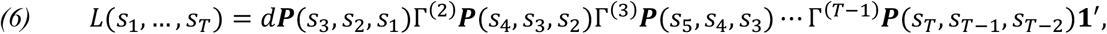

where *d* is the initial state distribution, ***P***(*s*_*i*_, *s*_*i*−1_, *s*_*i*−2_) is the *K* × *K* matrix

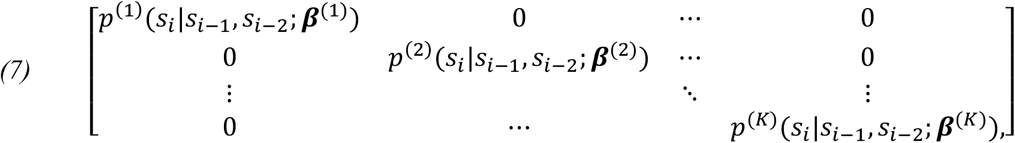

and **1**^′^ is the ones vector. As done by Klappstein et al. (2023), we maximize the likelihood directly using the forward recursive algorithm described by Langrock et al. (2012). Here we found that the BFGS optimization algorithm performs most consistently and provides the best estimates, in contrast to the Nedler-Mead method used as the default optimization algorithm in Klappstein et al. (2023). Regardless of the optimization method used, initial parameter values must be provided. The maximum likelihood estimates are sensitive to the initial parameter values, so we advise testing several sets of initial parameters when fitting the model. We recommend setting a burn-in period before fitting the data to the model to allow for the accumulation of memory. Confidence intervals for the parameters can be estimated when maximizing the likelihood directly via the approximate inverse Hessian matrix (Zucchini & MacDonald, 2009).

### 2.3 Simulation and validation

To assess the ability of our memory-based HMM-SSF model to estimate the weight that animals attribute to previous experiences when changing their habitat selection, we (1) simulated animal trajectories emerging from a state-dependent habitat selection process, with transition between states depending on the previous exposures to one or two covariates, and (2) fitted our memory-based HMM-SSF model to the simulated data to compare the estimated coefficients with the theoretical values used in the simulations. In the following, we present the results obtained with only one covariate, but similar results were observed when two covariates influenced the transition between states (Appendix B).

We used a two-state step selection function to simulate movement trajectories across simulated landscapes. For each set of parameters, we simulated 100 movement tracks, each of 5,000 time steps, in newly simulated landscapes (1000 × 1000 cells). The simulated trajectories emerged from two habitat selection patterns that differed in the strength of selection for a simulated habitat 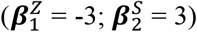 and in the strength of avoidance of movement cost 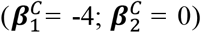. Another covariate (*R*) was influencing transition between states, with probabilities of transition to the second state increasing with higher covariate values 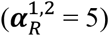 and, inversely, probabilities of transition to the first state decreasing with higher covariate values 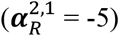. Habitats (*Z*) and (*R*) were randomly generated landscapes with values ranging between 0 and 1. The cost *C* was a nonlinear function of step length. These parameters were fixed across all simulations. The transition between states depended on the cumulative exposure to the covariate (*R*) rather than solely on the exposure at a given time. We used the coefficient (*δ*) to define the weight that simulated individuals attributed to previous experiences when deciding to change their habitat selection. Tested *δ* values ranged from low (0.1) to high (0.9) weights to mimic the importance given to short-term or long-term exposure to a covariate.

For each time step (*t*), 10,000 potential locations were generated from a uniform distribution of step lengths and turning angles around the location at time (*t*). The location at time (*t* + 1) was sampled among these potential locations with probabilities proportional to the state-dependent strength of selection. This strength of selection was determined by (1) the value of the habitat layer at the potential location and the cost to reach that location, (2) the state of the simulated individual at time *t*, and (3) the strength of attraction to the given habitat layer and the strength of avoidance of cost. At each time step, the exposure to the covariate (R) was extracted and used to update the internal state of the simulated individual following Equation [2].

We then fitted a memory-based HMM-SSF to each simulation. We used a 100-step burn in to allow for an initial accumulation of memory before fitting. For all values of the weight (*δ*) used in our simulations, past experiences that occurred more than 100 time steps ago were given a weight close to 0. Confidence intervals were estimated for all parameter estimates using the Hessian matrix (Zucchini & MacDonald, 2009). We also fitted a standard HMM-SSF to each simulation, using the *hmmSSF* function from the *hmmSSF* R package (Klappstein et al. 2023), to evaluate the bias in parameter estimates when accumulated experience is not included.

### 2.4 Application to Woodland Caribou

To illustrate the benefits of using a memory-based HMM-SSF, we analyzed a track of woodland caribou locations collected at 12-hour resolution from March-2011 to May-2013 in the boreal forest of Northern Ontario in Canada. Previous work showed that woodland caribou make movement decisions based on spatiotemporal variation in predation risk and resource availability (Avgar et al. 2015; McGreer et al. 2015; Viejou et al. 2018). We fitted a two-state HMM-SSF using our approach and compared the results to a standard HMM-SSF using the *hmmSSF* R package (Klappstein et al. 2023). In both cases, the SSF component captured selection coefficients for movement covariates (i.e., turning angles and step lengths), and the HMM component captured transition from area-restricted to exploratory movements as a function of the risk of predation and energetic gain. For this application, Equation [2] and Equation [3] therefore becomes

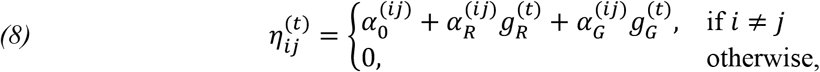

Where

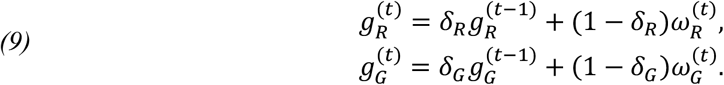

Here, 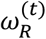 is the risk experienced at time *t* and 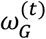 is the gain experienced at time *t*..

Spatiotemporal variation in the risk of predation and in energy availability have been previously evaluated in the study area and were available for this study (Avgar et al. 2015). Avgar et al. (2015) projected the spatial variation in dietary digestible biomass available to woodland caribou across the study area (spatial resolution: 500 meters) based on vegetation sampling to quantify forage abundance and digestibility and on animal-borne video observations to ascertain caribou diet (Thompson et al. 2015). Avgar et al. (2015) projected the spatial variation in predation risk across the study area (spatial resolution: 500 meters) based on radio-tracking data of 52 wolves from 34 different packs. For each caribou location, we extracted the values of wolf density and dietary digestible biomass and used them as a proxy on the perceived risk of predation and the energetic gain at a given time. Risk and energy values were centered and scaled, so our risk and energy maps reflected relative differences in risk of predation and energetic gain over space, rather than absolute values of risk and energetic gain. The model fitting was performed following the implementation methods described for the simulation study, testing 50 sets of initial parameters randomly chosen when fitting the model. Confidence intervals were computed for all parameter estimates via the approximate inverse Hessian matrix (Zucchini & MacDonald, 2009).

## 3 Results

### 3.1 Simulation study

Our memory-based HMM-SSF model adequately recovered the state-switching habitat selection parameters used in our simulations, both when estimating habitat selection coefficients, probabilities of transition between states, and the weights that simulated individuals attributed to previous experiences. In particular, weight coefficients were accurately estimated both when the simulated individuals relied primarily on previous experiences (*δ* = 0.9) or on current conditions (*δ* = 0.1) (Figure 1). Although the effect of a covariate on the transition probabilities between states was correctly estimated using our memory-based HMM-SSF, a standard HMM-SSF underestimated this effect (Figure 2). This bias was particularly strong when the simulated individuals relied primarily on their previous experiences to decide whether to change their habitat selection. Yet, habitat selection coefficients were correctly estimated from both our memory-based HMM-SSF and a standard HMM-SSF, both when simulated individuals relied on previous experiences or on current conditions when deciding to change their habitat selection (Appendix A. Figures S2, S3, S4, S5). Similar results were observed when transition between states depended on more than one covariate (Appendix B. Figures S6, S7).

**Figure 1.**
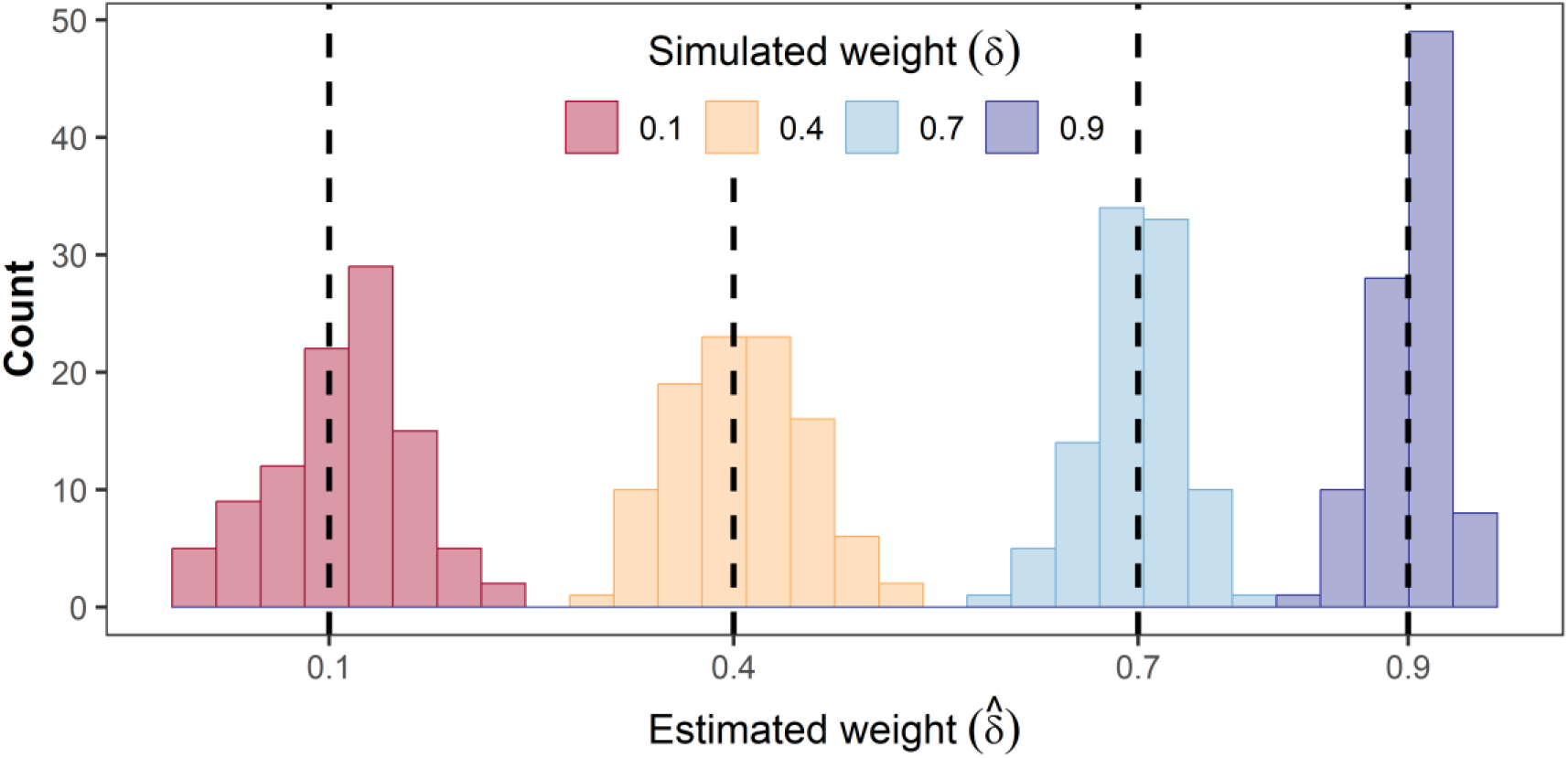
Distribution of the weights attributed to previous experiences as estimated using a memory-based HMM-SSF. Each distribution shows the estimated coefficients 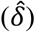 for 100 independent replications and is coloured according to the true weight (*δ*) used to simulate data (0.1, 0.4, 0.7, 0.9) and indicated by dotted vertical lines.

**Figure 2.**
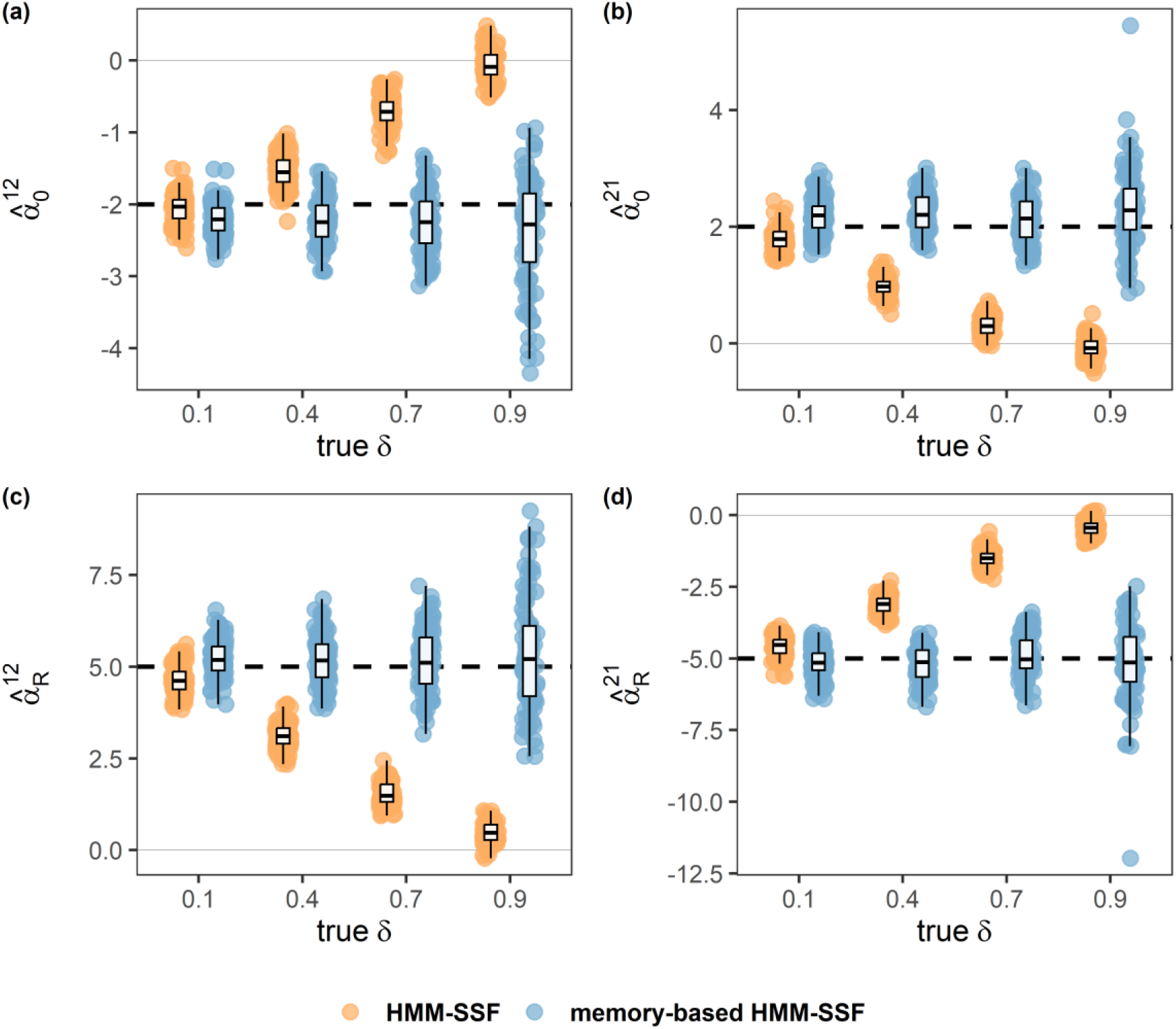
Comparison between state-transition coefficients estimated from a standard HMM-SSF (orange) and a memory-based HMM-SSF (blue). For each panel, the dotted line shows the true value of the state-transition coefficient used in the simulation. A higher delta coefficient indicates greater effect of an individual’s previous experience on transition between states. Each box represents the distribution of the estimated coefficients for 100 replications.

### 3.2 Application to woodland caribou

The monitored woodland caribou adopted area-restricted movements (*β*_*step length*_ = −10.8 [−12.4, −9.1]; *β*cos_(_*angle*_)_ = −0.10[−0.25, 0.05]) for an average of 11.1 days (*SE* = 3.4) before initiating an exploratory movement to a new area located, on average, 14.6 km away (*SE* = 1.8) (*β*_*step length*_ = −2.7 [−3.3, −2.1]; *β*cos_(_*angle*_)_ = 1.03[0.63, 1.44]) (Appendix A. Figure S6). The exploratory movements lasted 2.1 days on average (*SE* = 0.19), after which the animal returned to area-restricted movements. The probabilities of transitioning between area-restricted and exploratory movements depended on both long-term exposure to predation risk (*δ* = 0.96 [0.91, 0.98]) and short-term accumulated energetic gain (*δ* = 0.65 [0.28, 0.90]) (Figure 3). In particular, the probability of transitioning from exploratory to area-restricted movement increased significantly with higher accumulated energetic gain (*α*_*G*_ = 1.4 [0.21, 2.6]) and, inversely, decreased with higher exposure to predation risk (*α*_*R*_ = −2.7 [−4.9, −0.51]) (Figure 4). In contrast, these effects were not detected using a standard HMM-SSF (*α*_*G*_ = −0.21 [−1.2, 0.80]; *α*_*R*_ = 0.61 [−0.20, 1.4]), in which state transitions were modelled as functions of current predation risk and immediate energetic gain, rather than from exposure to risk accumulated over the last weeks or from energy accumulated over the last days.

**Figure 3.**
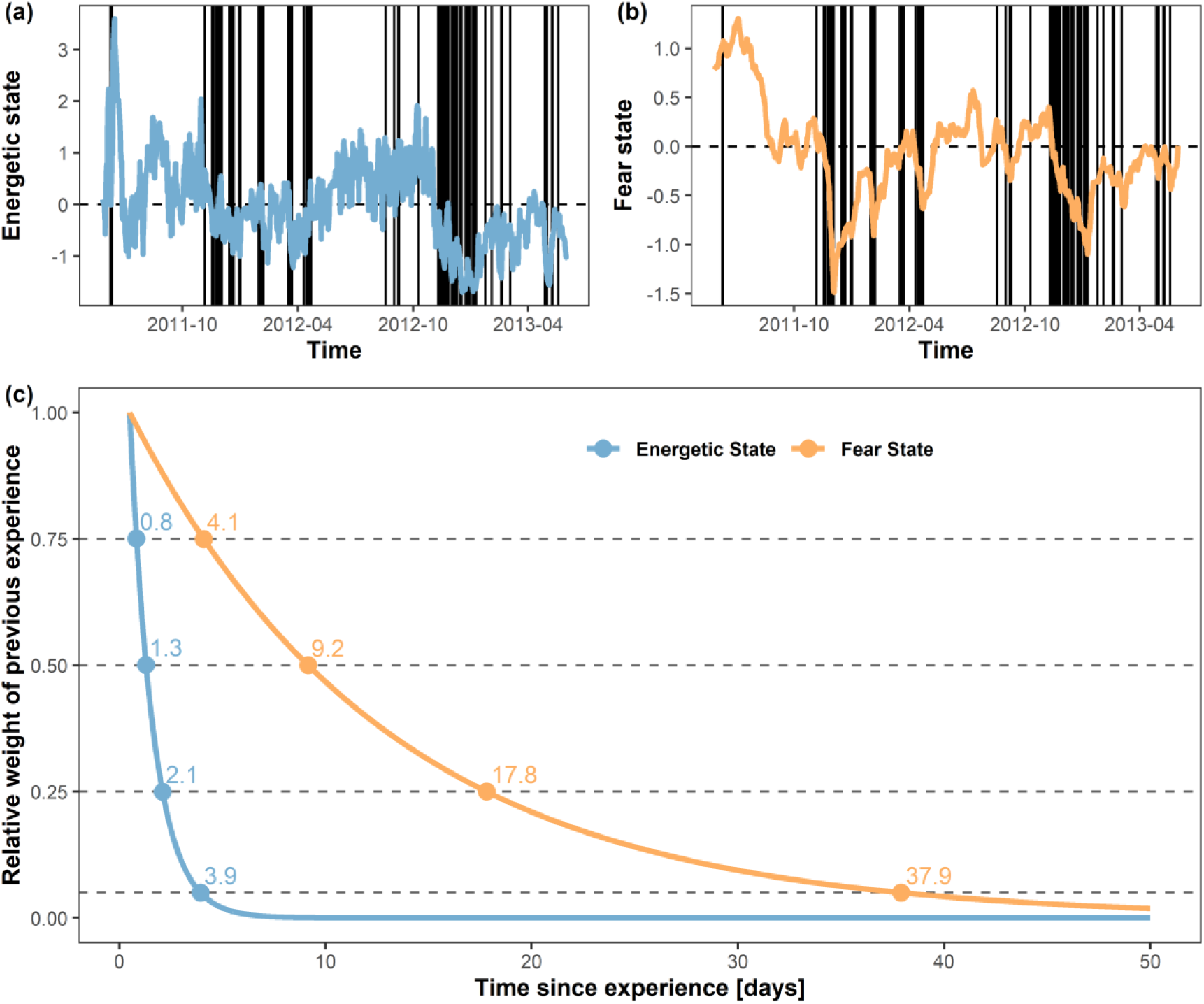
Dynamic internal states of a woodland caribou. Changes in the energetic (a) and fear (b) states are functions of the weight attributed to previous experiences (c). Transition between area-restricted and exploratory movements are indicated with black vertical lines in (a-b) and depended on long-term exposure to predation risk (weight = 0.96 [0.91 - 0.98]) and short-term accumulated energetic gain (weight = 0.65 [0.28 - 0.90]). The weight attributed to each previous experience is a monotonically decreasing function of time since the experience. For each time *t*, the weight of the experience at time (*t* − *ℓ*) is (1 − *δ*)*δ*^*t*−*ℓ*^. For graphical convenience, the y-axis shows the relative weight of the experience with respect to weight attributed to time t = 0.

**Figure 4.**
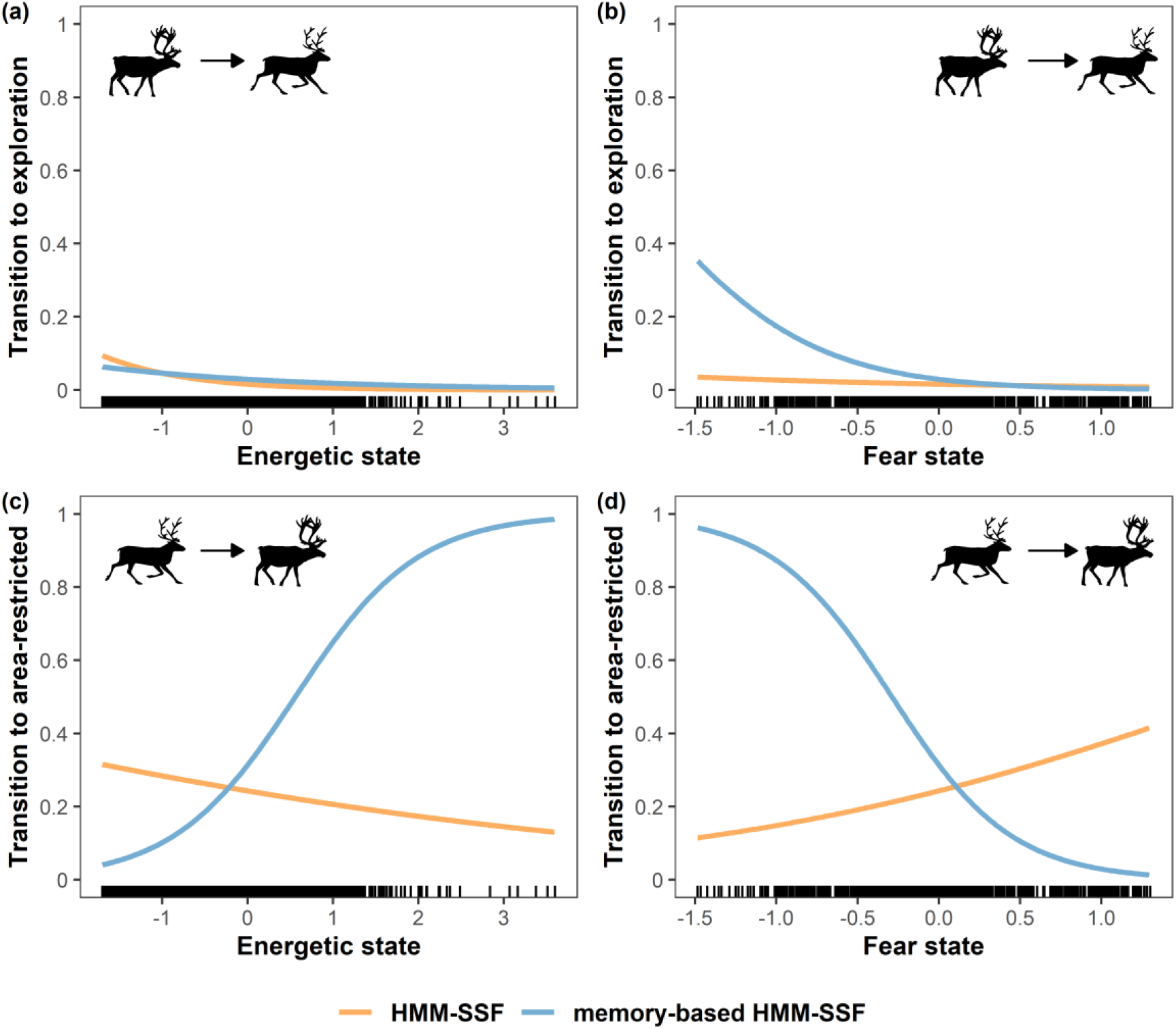
Probabilities of transition between area-restricted and exploratory movements as a function of energetic gain and predation risk. Effects of energetic gain and predation risk were estimated from a standard HMM-SSF (orange) and a memory-based HMM-SSF (blue). Energetic states accounted for the short-term accumulated energetic gain (weight = 0.65 [0.28 - 0.90]). Fear states accounted for the long-term exposure to predation risk (weight = 0.96 [0.91 - 0.98]).

## 4 Discussion

In this study, we introduce a memory-based state-switching habitat-selection model that explicitly accounts for animal experience when modelling movement decision-making. Our model extends the existing HMM-SSF (Klappstein et al. 2023; Pohle et al. 2024; Prima et al. 2022) by allowing changes in modes of habitat selection to depend on accumulated previous experiences that decay over time, rather than assuming that animals change their habitat selection decisions solely based on current external conditions. By integrating the accumulated experiences of animals into state dynamics, our memory-based state-switching habitat selection model explicitly incorporates changes in an animal’s internal state due to fitness-influencing processes, such as accumulated energy gain or accumulated risk of predation. Our generalized model therefore aligns with the movement ecology theoretical framework predicting that animal movement should be governed by external conditions as well as internal states (Nathan et al. 2008), allowing direct quantification of how temporally weighted past experiences influence state-dependent habitat selection.

### 4.1 A state-switching habitat selection model

Hidden Markov models are widely used in movement ecology to infer animal behaviour based on an animal’s motion pattern, especially from sequences of step length and turning angle (McClintock and Michelot 2018; Michelot et al. 2016). When coupled with step selection functions, HMM-SSF is an increasingly popular approach to infer behavioural-dependent habitat selection, such as quantifying habitat selection during area-restricted and exploratory behaviours (Klappstein et al. 2023; Pohle et al. 2024; Prima et al. 2022). However, it is important to clarify that HMM-SSF models fundamentally capture state-switching habitat selection patterns rather than only changes in movement patterns (i.e., based on an animal’s step length and turning angles). By capturing transition between modes of habitat selection, HMM-SSFs align with the recent development of time-varying resource-or step-selection functions to quantify temporal changes in habitat selection patterns (Dejeante et al. 2024; Klappstein et al. 2024). Augmenting the understanding provided by purely time-varying models, our memory-based state-switching habitat selection model applies a novel mechanistic framework to explain *why* animals change their mode of habitat selection by modelling the transition between states as a physiological or psychological process, i.e., as a function of dynamic internal conditions.

Importantly, our model ‘lets the data speak’, revealing whether animals rely on their internal state as accumulated through previous experiences to change their habitat selection over time or, alternatively, whether animals rely solely on current environmental conditions to decide where to move and when to change their mode of habitat selection. Critically, our simulation study shows that, when the true process underlying state-switching habitat selection depends on the long-term exposure to a given covariate, a standard HMM-SSF would underestimate the effect of this covariate, while our memory-based state-switching habitat selection model consistently recovers reliable estimates.

### 4.2 An experience-driven memory model

Animal movement decisions are rarely memoryless. Animals may remember the spatial areas they visited (spatial memory) or the environmental features associated to these locations (attribute memory) (Fagan et al. 2013). A rapidly growing body of evidence shows that animals rely on spatial or attribute memory to inform their movement decisions [for spatial memory see Dalziel et al. (2008); Falcón-Cortés et al. (2021); Merkle et al. (2019); Rheault et al. (2021); Schlägel et al. (2017); Thompson, Lewis, et al. (2022); for attribute memory see Avgar et al. (2015), (2013); Gurarie et al. (2022); Merkle et al. (2014); Ranc et al. (2021), (2022); Wolf et al. (2009)]. However, spatial and attribute memory are only one component of an animal’s internal memory. Cognitive-based movement models focus on quantifying where animals move in regard to space familiarity. Contrary to these existing models, our memory process models the impact of cumulative experiences on transition rates between behavioural states, evaluating why animals change their movement decisions based on previous experiences, rather than how animals select for memory-informed locations. Modelling animal memory as a temporally weighted sum of experiences offers new perspectives on evaluating the importance of physiological or psychological processes in movement ecology. Future studies could integrate our memory-based movement model with cognitive maps to jointly account for memory-driven motivational states and spatial or attribute memory. This would make it possible to quantify not only where animals move in regard to space familiarity, but also how accumulated exposure to environmental variables (such as risk of predation or energy) influence the behavioural movement states. For example, one could test whether individuals are more likely to select areas with known forage quality when they have accumulated high levels of energetic stress (i.e., attribute memory) or whether individuals are more likely to select familiar places when they have accumulated high levels of predation risk (i.e., spatial memory).

### 4.3 A dynamic state of fear

By estimating the weight attributed to previous experiences, our memory-based state-switching habitat selection model can determine whether animals integrate predation risk experienced over the short-or long-term when deciding to change their habitat selection. Prey individuals can adopt proactive and reactive responses to reduce the risk of predation, responding to the immediate perception of a predator (Broekhuis et al. 2013; Martin and Owen-Smith 2016) or to long-term exposure to predation risk (Creel 2018; Valeix et al. 2009). Proactive responses are often based on learning or memory processes and require long-term, predictable exposure to the risk of predation (Creel 2018). Prey individuals can, for example, avoid areas associated with higher intensity of use by a predator (Valeix et al. 2009). Alternatively, places perceived as riskier could reflect prey recognition of the habitat types favoured by their predators (Avgar et al. 2015; McGreer et al. 2015). Animals therefore move through a dynamic landscape of fear which characterizes the spatiotemporal variation in the risk of predation that prey perceive, leading to the adoption of anti-predation behavioural strategies (Laundre et al. 2010; Palmer et al. 2022).

In addition to remembering the spatial areas associated with a higher predation risk (i.e., attribute memory), cognitive processes may determine the level of fear of an individual, which may depend on the accumulated exposure to a risk of predation rather than on the immediate perception of such risk. For example, our application shows that a woodland caribou can use information of predation risk acquired up to 15 days earlier to decide whether to settle in a given area or whether to continue exploring its landscape. By incorporating past experiences into the internal dynamics that govern transitions between states, our framework allows one to infer the “dynamic state of fear”, i.e., the memory-weighted accumulation of perceived predation risk driving changes in animal movement decisions.

### 4.4 A dynamic energetic state

Although animals may respond to long-term exposure to the perceived risk of predation, changes in habitat selection may also result from short-term exposure to specific environmental conditions. Theoretical studies show that animal behavioural decisions should depend on physiological states, such as energetic states (Houston et al. 1993; McNamara and Houston 1992). There is empirical evidence that gut fullness or hunger state can mediate transitions between area-restricted and exploratory movements, with individuals adopting exploratory movement when hungry and area-restricted searches once a rich feeding patch is found (Dorfman et al. 2022; Kareiva and Odell 1987). This mixture of random walks is recognized as an efficient foraging strategy (Benhamou 1992, 2007). Contrary to existing HMM-SSF models, a memory-based state-switching habitat selection model allows one to model such a movement pattern by explicitly accounting for the dynamic energetic state that governs these changes between exploratory and area-restricted modes of movement.

For example, our application to woodland caribou illustrates that an individual is more likely to settle in a given space when the accumulated energetic gain has been elevated over the last 2 days, a signal that is underestimated when only considering the amount of energy available at a given time. Integrating an animal’s previous experiences into state-switching habitat selection models therefore allows one to explicitly consider the dynamics of energetic state, similarly to continuous-time recharge-dynamic movement models (Hooten et al. 2019).

## 5 Conclusion

Our memory-based state-switching habitat selection model provides an accessible yet powerful framework for investigating how animal experiences influence state-switching habitat selection. This framework enables dynamic internal states to be modelled as functions of accumulated physiological or psychological conditions, extending the role of memory beyond spatial familiarity to include memory of internal states. By integrating memory and physiology within a single inferential framework, our memory-based state-switching habitat selection model broadens the scope of movement ecology from explaining not only where animals move (i.e., habitat selection) but also why they move (i.e., changing their habitat selection in response to motivational states). For example, state-switching movement decisions can be modelled as a function of an animal’s energetic condition, stress, or anxiety state, providing new opportunities to link movement behaviour with underlying physiological and psychological processes.

### Statement of authorship

RD and AK led the study and model design, with substantial contributions from JMF and MAL. JMF provided data for the caribou application of the model. All authors contributed conceptual ideas throughout the project. RD and AK conducted the analyses and led the manuscript writing, with contributions from all co-authors.

## Acknowledgments

Financial support for this research was provided by the Forest Ecosystem Science Cooperative, Ontario Ministry of Natural Resources and Forestry, the National Science and Engineering Council of Canada [funding reference number CGSD - 600060 – 2025], The Ontario Ministry of the Environment, Conservation and Parks, and Canadian Forest Service (CFS) Policy Branch.

## Competing interests

The authors declare that they have no conflict of interest.

## Appendix

## A. Supplemental figures

**Figure S1.**
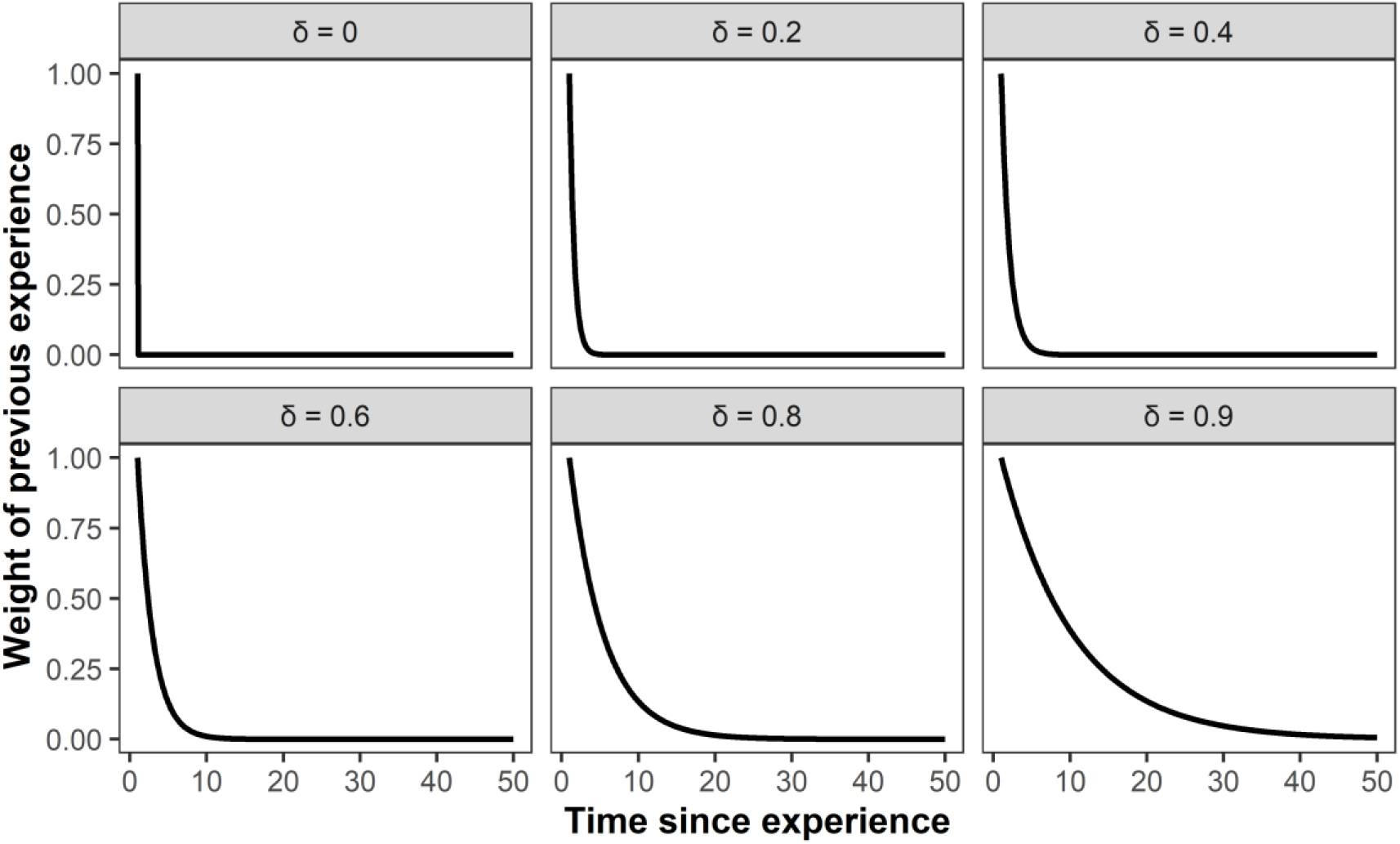
Weighting function of previous experiences based on (*δ*) values. The weight attributed to each previous experience is a monotonically decreasing function of time since the experience. For each time *t*, the weight of the experience at time (*t* − *ℓ*) is (1 − *δ*)*δ*^*t*−*ℓ*^. For graphical convenience, the y-axis shows the relative weight of the experience with respect to weight attributed to time t = 0.

**Figure S2.**
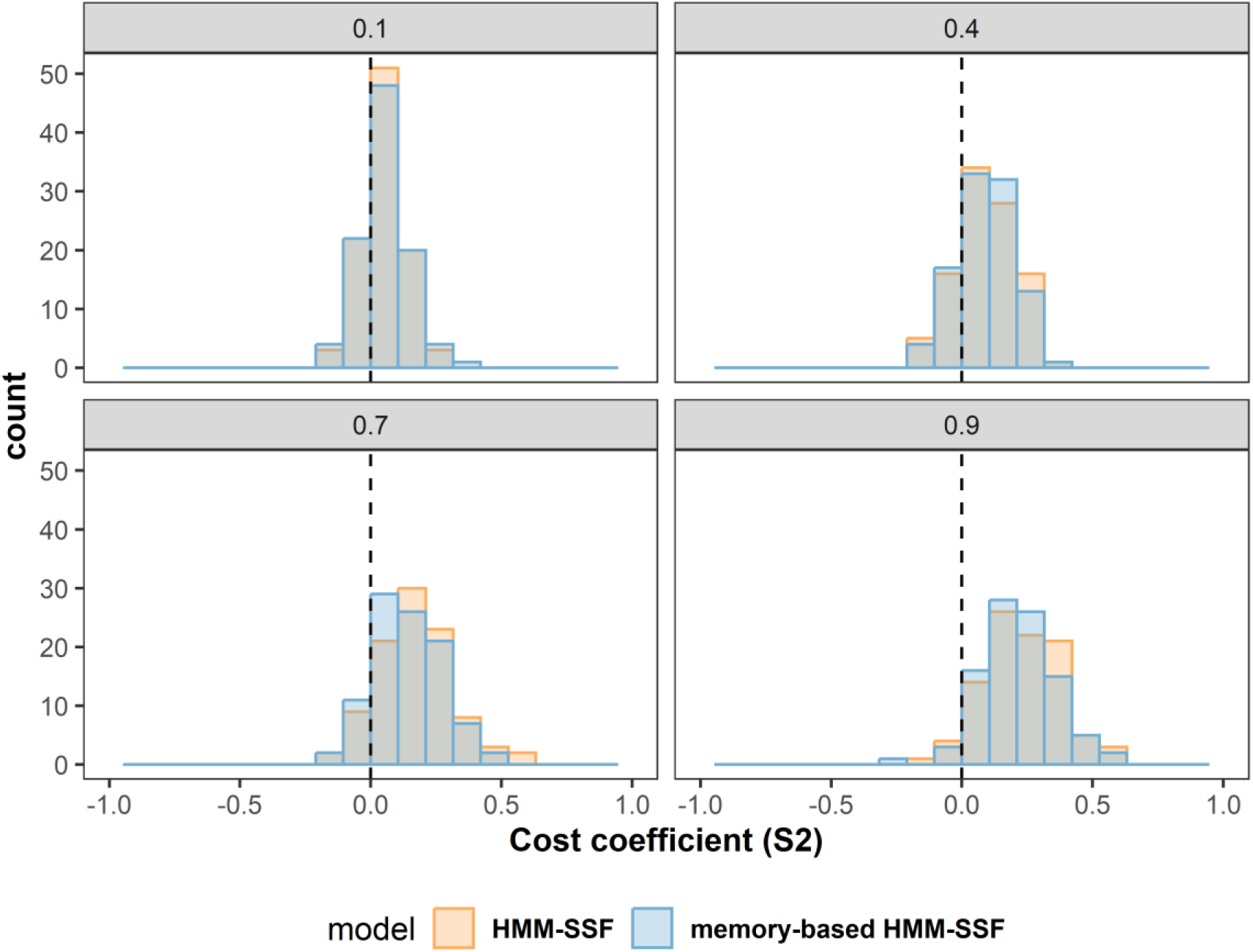
Distribution of the estimated coefficients of avoidance of cost in state 2. The dotted vertical line shows the true coefficient value used in the simulation. Coefficients were estimated using a standard HMM-SSF (orange) or using a memory-based HMM-SSF (blue).

**Figure S3.**
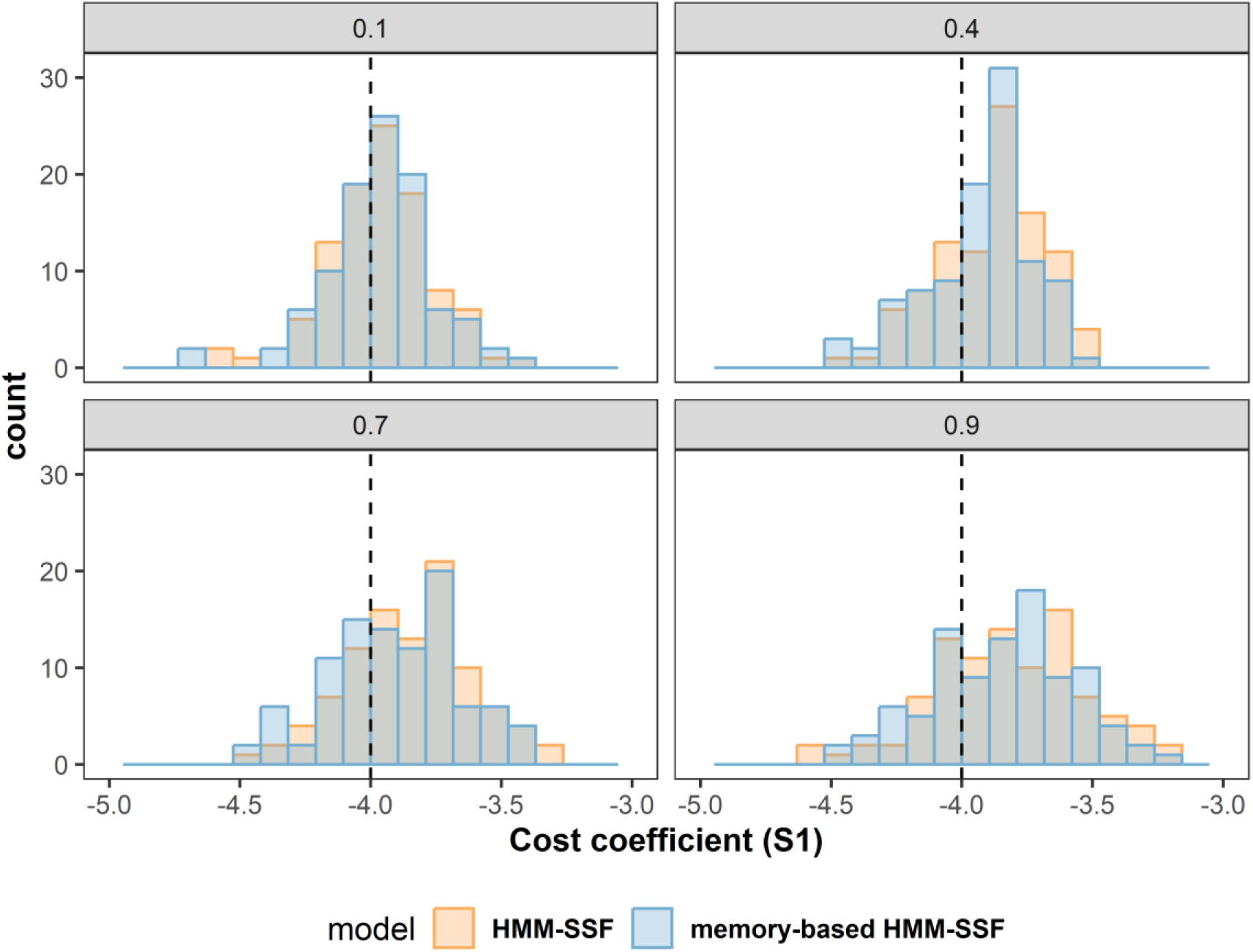
Distribution of the estimated coefficients of avoidance of cost in state 1. The dotted vertical line shows the true coefficient value used in the simulation. Coefficients were estimated using a standard HMM-SSF (orange) or using a memory-based HMM-SSF (blue).

**Figure S4.**
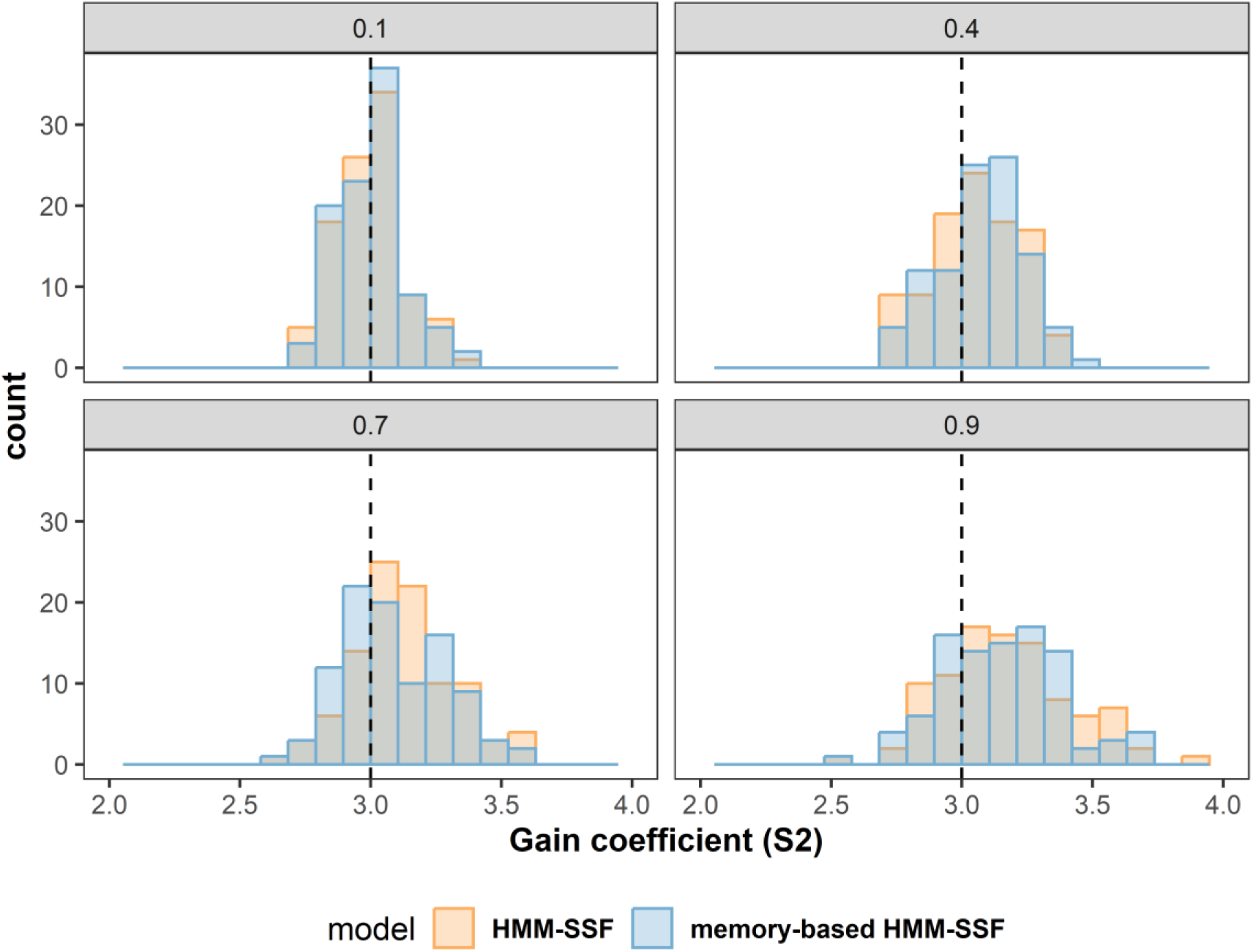
Distribution of the estimated coefficients of avoidance of gain habitat in state 2. The dotted vertical line shows the true coefficient value used in the simulation. Coefficients were estimated using a standard HMM-SSF (orange) or using a memory-based HMM-SSF (blue).

**Figure S5.**
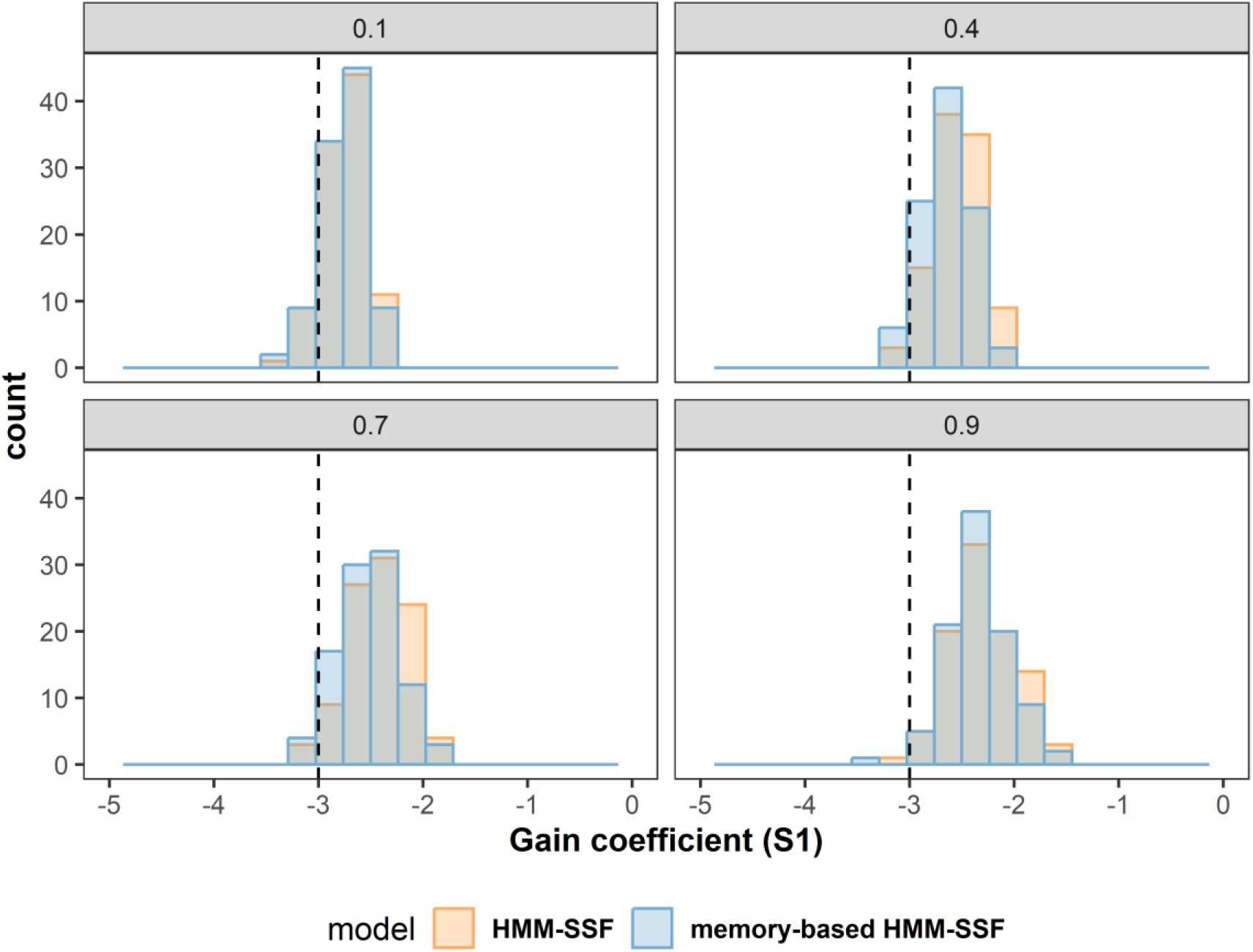
Distribution of the estimated coefficients of avoidance of gain habitat in state 1. The dotted vertical line shows the true coefficient value used in the simulation. Coefficients were estimated using a standard HMM-SSF (orange) or using a memory-based HMM-SSF (blue).

**Figure S6.**
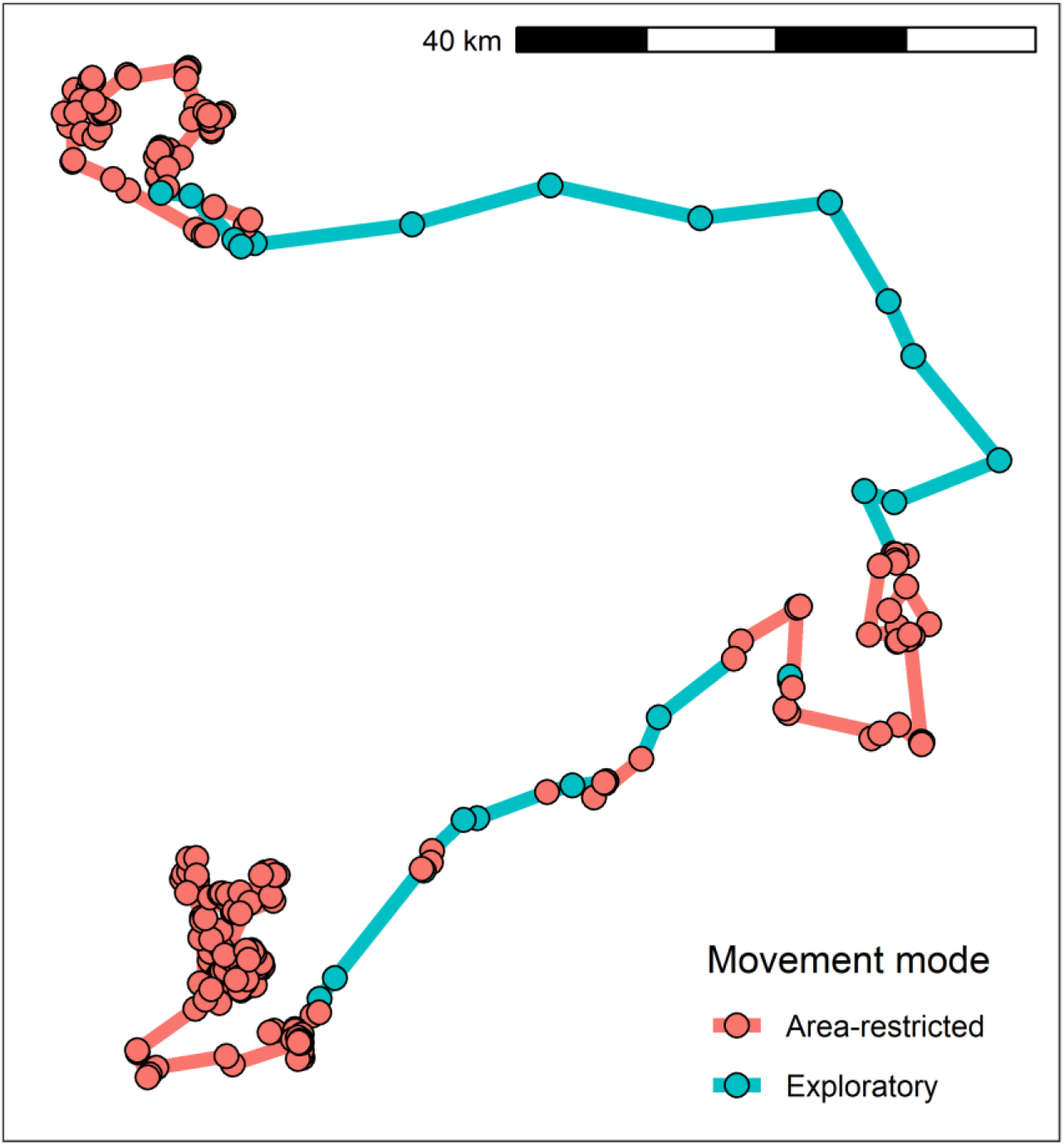
Movement trajectory of the studied woodland caribou obtained from 2012-02-06 to 2012-08-06 with locations recorded every 12h30. Movement modes are inferred from the memory-based HMM-SSF described in section 2.4.

## B. Simulation details with two covariate influencing transition between states

For the simulation study, we define the step selection component to be

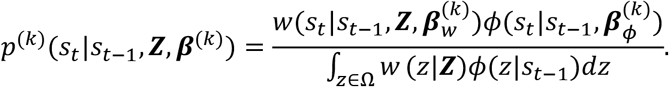

The weighting equation is

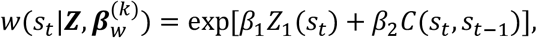

where *Z*_1_ is the habitat covariate and *C* is the cost function given by

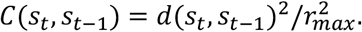

Here, *d* is the Euclidean distance between *s*_*t*_ and *s*_*t*−1_ (i.e. the step length) and *r*_*max*_ is the maximum theoretical distance an animal could travel in one time step, fixed *a priori* for both the simulation and the fitting. The movement kernel Φ is the product of a uniform distribution of step lengths bounded by a circle of radius *r*_*max*_ and turning angles.

The transition matrix for the HMM is

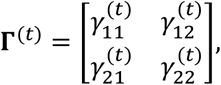

with

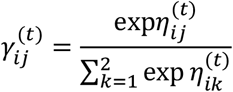

and

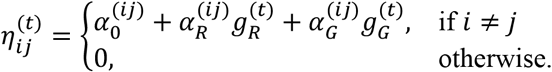

for the two covariate model. We define

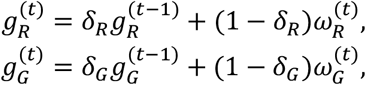

where 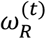 is the risk experienced at time *t* and 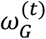 is the gain experienced at time *t*.

The parameters for the simulations are described in Table 1. These parameters remained fixed for all simulation runs. When using two covariates in the HMM, we tested five unique combinations of *δ*_*R*_ and *δ*_*G*_ to ensure these parameters could be reliably recovered from movement data. For each combination, we generated 100 simulations of 5,000 steps.

**Table S1.** Parameters for simulation model with two covariates influencing transition between states.

| Description | Parameter | Value |
| --- | --- | --- |
| Intercept | $\alpha_0^{(12)}$ | -2 |
| Intercept | $\alpha_0^{(21)}$ | -2 |
| Effect of risk in transition from state 1 to 2 | $\alpha_R^{(12)}$ | 2 |
| Effect of risk in transition from state 2 to 1 | $\alpha_R^{(21)}$ | -2 |
| Effect of gain in transition from state 1 to 2 | $\alpha_G^{(12)}$ | -5 |
| Effect of gain in transition from state 2 to 1 | $\alpha_G^{(21)}$ | 1 |
| Effect of habitat on step selection in state 1 | $\beta_1^{(1)}$ | -3 |
| Effect of habitat on step selection in state 2 | $\beta_1^{(2)}$ | 3 |
| Effect of cost on step selection in state 1 | $\beta_2^{(1)}$ | -5 |
| Effect of cost on step selection in state 2 | $\beta_2^{(2)}$ | -2 |
| Weight attributed to previous experiences of risk | $\delta_R$ | 0.1, 0.4, 0.8 |
| Weight attributed to previous experiences of gain | $\delta_G$ | 0.1, 0.4, 0.8 |

**Figure S7.**
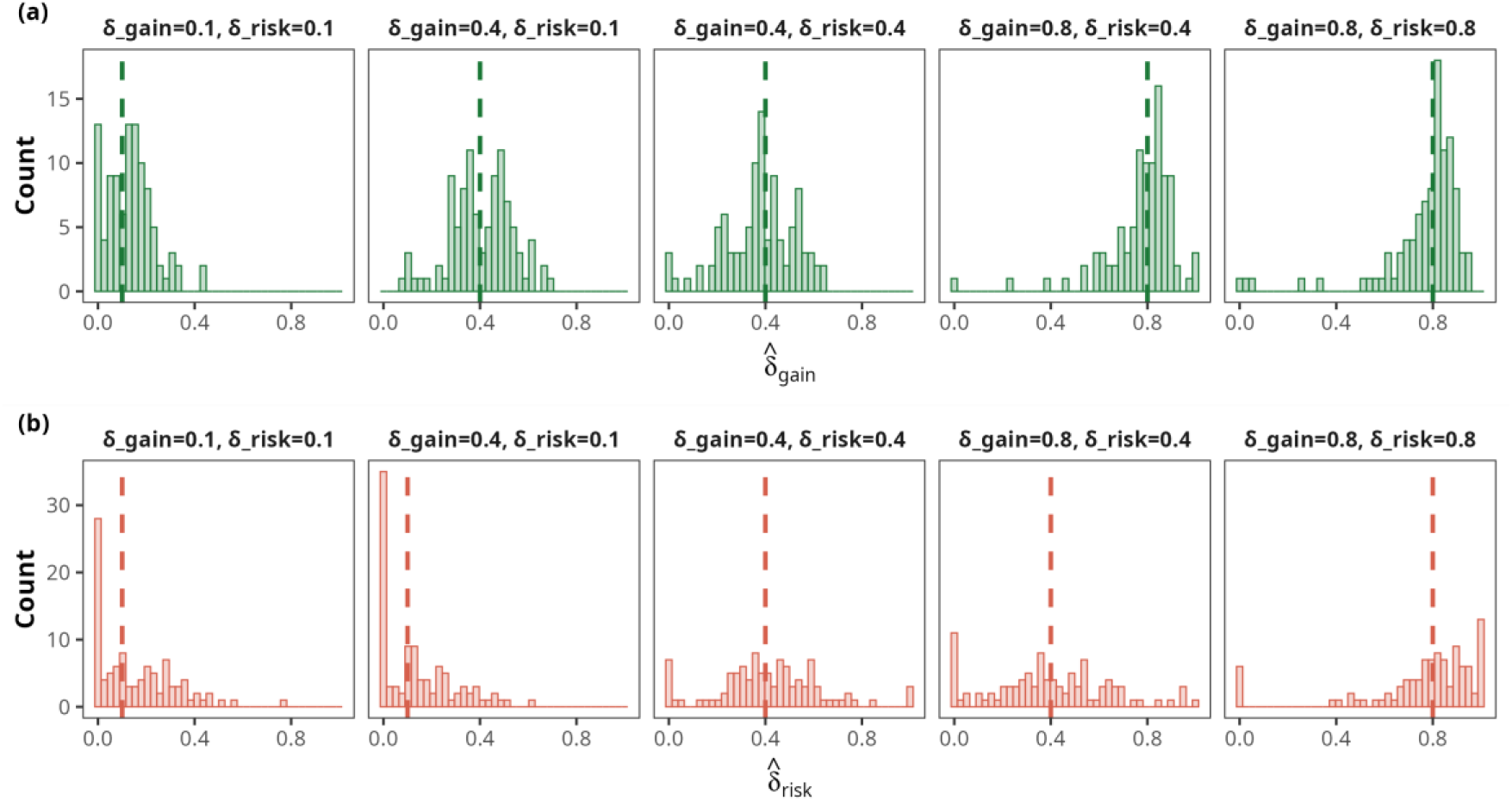
Distribution of the weights attributed to previous experiences as estimated using a memory-based HMM-SSF. Each distribution shows the estimated coefficient pairs 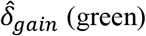 or 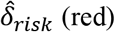 for 100 independent replications and is labelled according to the true weights (*δ*_*risk*_, *δ*_*gain*_) used to simulate data and indicated by dotted vertical lines.

**Figure S8.**
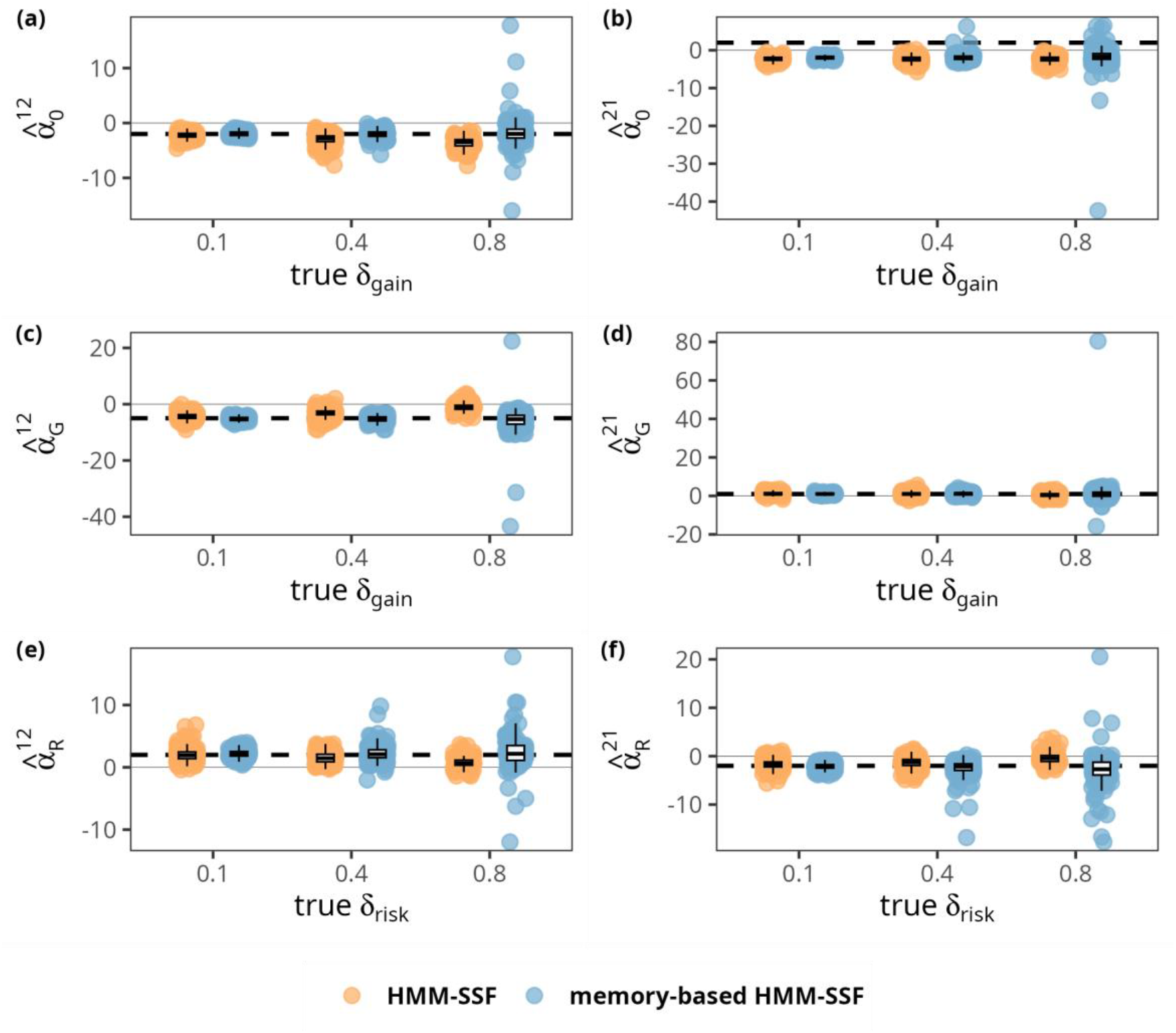
Comparison between state-transition coefficients estimated from a standard HMM-SSF (orange) and a memory-based HMM-SSF (blue). For each panel, the dotted line shows the true value of the state-transition coefficient used in the simulation. A higher delta coefficient indicates greater effect of an individual’s previous experience on transition between states. Each box represents the distribution of the estimated coefficients for 100 replications.

## Notes

### Competing Interest Statement

The authors have declared no competing interest.

